# Gut mucosal cells transfer α-synuclein to the vagus nerve

**DOI:** 10.1101/2023.08.14.553305

**Authors:** Rashmi Chandra, Arpine Sokratian, Katherine R. Chavez, Stephanie King, Sandip M. Swain, Joshua C. Snyder, Andrew B. West, Rodger A. Liddle

**Author notes:** Denotes equal contributions (Rashmi Chandra, Arpine Sokratian). **Co-Corresponding authors:** Rodger A. Liddle, M.D. Box 103859 1033A Genome Science Research Building -1 905 LaSalle Street Duke University Medical Center Durham, NC 27710 Andrew B. West, Ph.D. 3 Genome Ct., Durham, NC, 27710.

## Abstract

Epidemiological and histopathological findings have raised the possibility that misfolded α-synuclein protein might spread from the gut to the brain and increase the risk of Parkinson’s disease (PD). While past experimental studies in mouse models have relied on gut injections of exogenous recombinant α-synuclein fibrils to study gut to brain α-synuclein transfer, the possible origins of misfolded α-synuclein within the gut have remained elusive. We recently demonstrated that sensory cells of the gut mucosa express α-synuclein. In this study, we employed mouse intestinal organoids expressing human α-synuclein to observe the transfer of α-synuclein protein from gut epithelial cells in organoids co-cultured with vagal nodose neurons that are otherwise devoid of α-synuclein expression. In intact mice that express pathological human α-synuclein, but no mouse α-synuclein, α-synuclein fibril templating activity emerges in α-synuclein seeded fibril aggregation assays in tissues from the gut, vagus nerve, and dorsal motor nucleus. In newly engineered transgenic mice that restrict pathological human α-synuclein expression to intestinal epithelial cells, α-synuclein fibril-templating activity transfers to the vagus nerve and to the dorsal motor nucleus. Subdiaphragmatic vagotomy prior to the induction of α-synuclein expression in the gut epithelial cells effectively protects the hindbrain from the emergence of α-synuclein fibril templating activity. Overall, these findings highlight a novel potential non-neuronal source of fibrillar α-synuclein protein that might arise in gut mucosal cells.

## Introduction

Parkinson’s disease (PD) is a debilitating neurodegenerative disease with characteristic motor disturbances including rigidity, resting tremor, and bradykinesia. Many patients also suffer from gastrointestinal symptoms such as constipation that often precede characteristic motor deficits by ten years or more ^1^. The pathological hallmarks of PD are intracellular proteinaceous inclusions filled with fibrillated forms of α-synuclein that accumulate in both the brain and peripheral nervous system. In dopaminergic neurons in PD, inclusions known as Lewy bodies have been associated with neuronal vulnerability and degeneration ^2,3^. Throughout the brain, α-synuclein is normally found in presynaptic terminals of excitatory neurons and other neuronal subtypes, with a role in endocytosis and synaptic vesicle function ^4^. Mutations in the α-synuclein gene (SNCA) such as A53T and A30P, as well as multiplication of the SNCA locus, can cause familial PD ^5,6^. One of the more remarkable features of α-synuclein protein is the intrinsic ability to aggregate into β-sheet rich protein fibrils that have high affinity towards amyloid dyes like thioflavin ^7–9^. These α-synuclein fibrils have a proposed capacity to spread between inter-connected cells in an hypothesized prion-like cascade ^10–13^. Transferred α-synuclein might recruit native α-synuclein within the recipient cell to seed additional aggregates ^14–16^ that can form larger fibrils and inclusions^17,18^. The presence of α-synuclein fibrils and templating activity in the brain and cerebrospinal fluid has been convincingly demonstrated in PD as measured through newly developed seeded aggregation assays (SAAs) that include protein misfolding cyclic amplification (PMCA) and real-time quaking-induced conversion (RT-QuIC) assays ^19^. The α-synuclein RT-QuIC assay demonstrated seeding activity in duodenal biopsies of PD patients but not in healthy control subjects ^20^. The originating source of the α-synuclein seeds that trigger activity in this assay are not clear. In rat models, it is thought that seeds that trigger pathological accumulations of α-synuclein may orginate in neurons and the brain and descend into the gut or, originate somewhere in the gut and ascend into the brain ^21^.

Clinical and experimental data indicate that the gut may play a role in PD susceptibility. Not only do gastrointestinal symptoms such as constipation often precede the motor symptoms of PD ^22–24^, experimentally, it has been suggested in rodent models that exogenous α-synuclein fibrils introduced into the gut can spread to the brain ^25–27^. Abnormal α-synuclein aggregates have been histopathologically identified in the enteric nervous system prior to the development of PD ^22^. Evidence supporting a role for the enteric nervous system involvement in α-synuclein pathology includes the observation that α-synuclein immunoreactive inclusions localize to neurons of the submucosal plexus, whose axons project to the gut mucosa ^28–30^ and myenteric plexus, and parallels input of the vagus nerve ^31^. Mice harboring the A53T transgene, exhibited enteric nervous system dysfunction including increased colonic transit times and reduced fecal output consistent with constipation ^32,33^. In animals, the vagal route of α-synuclein transport has also been documented following exposure to the environmental toxicant rotenone that can cause α-synuclein misfolding ^34^, as well as direct injections of adeno-associated viral vectors overexpressing human α-synuclein in vagal neurons ^35^. More recently, it was demonstrated that vagotomy prevented the formation of brain aggregates when α-synuclein fibrils were injected into the gut of susceptible mice, indicating that pathogenic fibrils can spread from nerve terminals in the gut to the brain^25,36^. However, the physiologically relevant origins of misfolded α-synuclein protein that might use the vagus nerve as a conduit into (or out of) the brain have been unclear. One clue might be that some epidemiological observations suggest that complete truncal vagotomy in patients is associated with a decreased risk of PD, suggesting in at least some PD cases that α-synuclein aggregation in the gut may be critical for PD risk later on ^37–39^.

Recently, it was discovered that enteroendocrine cells (EECs) in the gut mucosa connect with neurons in culture and in the intact gut of mice ^40^. Using modified rabies viral tracing in vivo, a synaptic connection was confirmed between PYY-expressing EECs and neurons in the colon ^40^. In co-culture experiments in dishes, cholecystokinin (CCK)-containing EECs and sensory neurons form spontaneous synaptic connections ^40,41^. Thus, there appears to be an inherent affinity between EECs and neurons ^42^. EECs are exposed to the lumen of the gut and respond to numerous chemical and physical stimuli from diet, the microbiome, and the environment ^40,43–45^. Long thought to produce exclusively gut hormones, it is now known that EECs possess neuron-like properties, and part of that neuronal-like phenotype might include the endogenous expression of α-synuclein ^42,46^. EECs express neurotransmitters and other canonical pre-synaptic proteins, and possess axon-like processes sometimes called neuropods through which they connect to nearby nerves ^40,41^. To explore a potential novel cellular source of pathological α-synuclein that might contribute seeding activity from the gut to the brain, here we use organoids and transgenic mice to explore whether α-synuclein has a potential to spread from gut mucosal cells to the vagus nerve.

## Results

In the gut, EECs present as elongated or flask-shaped cells and express high levels of α-synuclein protein according to our past immunohistochemical analyses ^46^. The apical surface of EECs is typically open to the intestinal lumen, and the basal surface lies on the lamina near neurons (Fig. 1A). To study the possible spread of pathological (i.e., human A53T α-synuclein) α-synuclein protein, we established a humanized α-synuclein mouse model, in which endogenous Snca was deleted and instead replaced with α-synuclein protein expressed from a PAC encoding human A53T α-synuclein (SNCA^A53T^). This mouse strain is known to develop mild but widespread pS129-α-synuclein protein accumulations with physiological levels of α-synuclein levels ^32,47^, and develops α-synuclein immunoreactive aggregates in enteric ganglia by three months of age as well as early gastrointestinal dysfunction ^32^. Cck-eGFP mice (which express enhanced green fluorescent protein in EECs), also devoid of endogenous mouse α-synuclein, were crossed with mice expressing the PAC-Tg(SNCA^A53T^) ^48^ to generate PAC-Tg(SNCA^A53T^;Cck-eGFP;Snca^-/-^). This mouse line is referred to hereafter as SNCA^A53T^. SNCA^A53T^ mice develop normally and exhibit human α-synuclein expression that localizes primarily to the basal pole of EECs in the gut (Fig. 1B). Cck-eGFP expression demarcates some EECs in the proximal intestine ^40,41^. Vagal innervation of the gut is most dense in the proximal intestine and diminishes in the colon ^49^, therefore, to investigate the potential neuronal spread of α-synuclein we focused on the small intestine. According to sensitive ELISA analysis for human α-synuclein protein, α-synuclein expression in the vagal nodose ganglia was low (∼170 pg per mg of tissue, Fig. 1C) compared to much higher expression in the duodenum and upper intestine (∼140 ng per mg of tissue, Fig. 1D), with lesser expression in the hindbrain that encompasses the dorsal motor nucleus (DMV, Fig. 1E). Despite the much lower levels of α-synuclein protein in nodose ganglia compared to the hindbrain, the isolated ganglia tissue harbored positivity for fibril-seeding activity in RT-QuIC assays (dwell time to threshold 5.79 hrs / 1.43 hours, Fig. 1F). Dwell times in RT-QuIC reactions indicated possible femtomolar concentrations of α-synuclein fibril seeds in the lysates from the nodose tissue according to standard curves of recombinant fibrils of known concentrations (Supplemental Fig. 1). Comparable RT-QuIC signals in gut and hindbrain tissues were also observed corresponding to dwell times to threshold of 19.1 hours / 5.5 hours (Fig. 1G,H). Matched tissues dissected and processed at the same time from the same region in the mice that lacked human α-synuclein expression did not reveal any comparible fibril templating activity (Fig. 1F-H). These results extend the observations of Kuo et al., who originally demonstrated the presence of enteric α-synuclein pathology with immunohistochemistry in the gut in the PAC-Tg(SNCA^A53T^) mouse strain ^32^.

**Figure 1.**
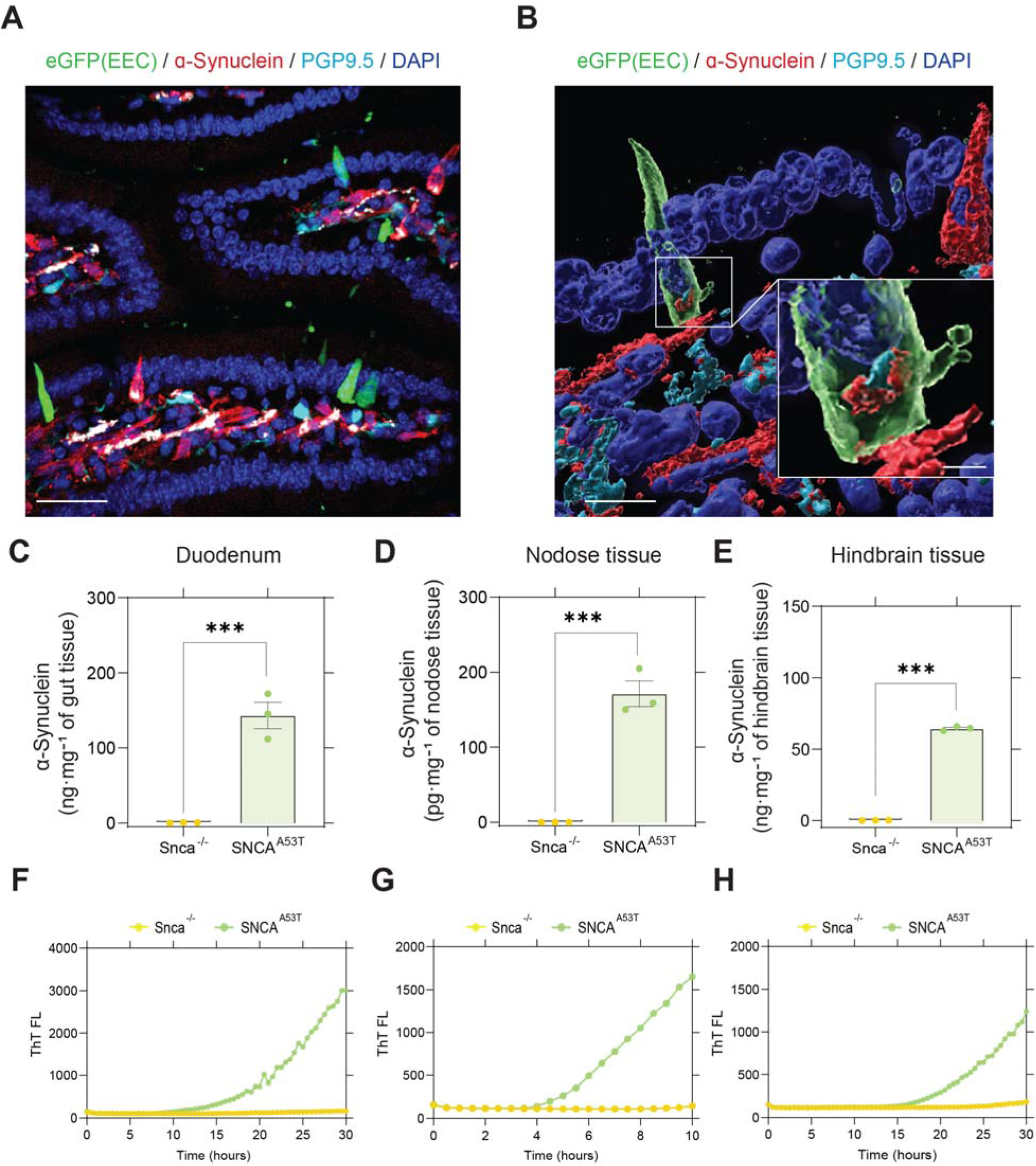
Alpha-synuclein expression and seeding activity in SNCA^A53T^ mice. (A,B) Immunostaining of duodenum harvested from SNCA^A53T^ mice. Enteroendocrine cells (EECs) expressing green fluorescent protein (green) are scattered among other mucosal cells (DAPI labeled nuclei, blue) and are in proximity to α-synuclein (red) containing fibers stained with the pan-neuronal marker PGP9.5 (cyan) in the lamina propria of the villus. ELISA quantification of human α-synuclein in (C) duodenum (α-synuclein quantification in nanograms per mg of nodose tissue), (D) nodose ganglia (α-synuclein quantification in picograms per mg of nodose tissue), and (E) hindbrain (α-synuclein quantification in nanograms per mg of nodose tissue), from Snca^-/-^ and SNCA^A53T^ mice. RT-QuIC analysis of (F) duodenum, (G) nodose ganglia, and (H) hindbrain of Snca^-/-^ and SNCA^A53T^ mice. Scale bars are 30 μm in panel A, 10 μm and 1 μm (inset) in panel B. Significance was determined by unpaired t-test, where ***P<0.001.

Recognizing that pathological α-synuclein is expressed in EECs in the model, and EECs lie in close proximity to enteric submucosal neuronal fibers, we next sought to determine if α-synuclein expressed in EECs might spread to adjacent nerves. To evaluate this possibility, we co-cultured intestinal organoids from the SNCA^A53T^ mice with nodose ganglia neurons from Snca^-/-^ mice that lack any α-synuclein expression (Fig. 2). Though enteric ganglia might natively express low levels of α-synuclein protein (Fig. 1C), culturing the neurons from the Snca^-/-^ mice and utilizing a monoclonal anti-human α-synuclein antibody previously validated in the Snca^-/-^ mice ^50^ ensures reliability for detection of authentic α-synuclein spread from the intestinal organoids to the co-cultured nerves. In the organoids, the EECs (identified by eGFP expression) are oriented with their apical surface open to the lumen (Fig. 2B). Given that organoids remain stationary within the Matrigel matrix, it appeared that nerve fibers with strong PGP9.5 or neuron-specific class III tubulin (Tuj1) expression from the Snca^-/-^ mice grew towards the basal surface of EECs, demonstrating that organoids attract and connect with nerve fibers in vitro. Such results are consistent with previous observations with isolated EECs^40,41,51^. As expected, neuronal fibers from Snca^-/-^ mice exhibited no detectable α-synuclein via immunohistochemistry (Supplemental Fig. 2). Within five days, some nerve fibers had grown towards organoids and appeared to be in close contact with EECs (i.e., CCK-positive cells, Fig. 2C-E). CCK-positive EECs from SNCA^A53T^ mice (green) in intestinal organoids express readily detectable α-synuclein (red). More striking was the stark appearance of α-synuclein protein within PGP9.5 (a pan-neuronal-specific marker) neuritic outgrowths nearby CCK-eGFP positive cells (Fig. 2C). While α-synuclein protein distributed in the cytoplasm of EECs, α-synuclein protein co-localized within neurites from Snca^-/-^ mice in patches along the process, presented as patches along the neuritic PGP9.5 positive length. Transferred α-synuclein protein could be detected along PGP9.5 processes on the basal organoid surface running next to EEC cells (Fig. 2C). Alpha-synuclein protein was similarly observed in Tuj1 positive neurites in which neuronal fiber staining appeared more uniform (Fig. 2D-E). Because of the high densities inherent to the organoid systems, and high connectivity and dynamic outgrowth of co-cultured ganglia, it was not possible to ascertain a critical distance that facilitated the transfer of α-synuclein onto adjacent neuronal fibers even though other studies in 2D chambered cells suggest direct cell-to-cell contact is required for transfer^42^. Alpha-synuclein protein appeared to emerge from basal and lateral regions of the EECs that were also in physical contact with PGP9.5-or Tuj1-positive ganglionic outgrowths.

**Figure 2.**
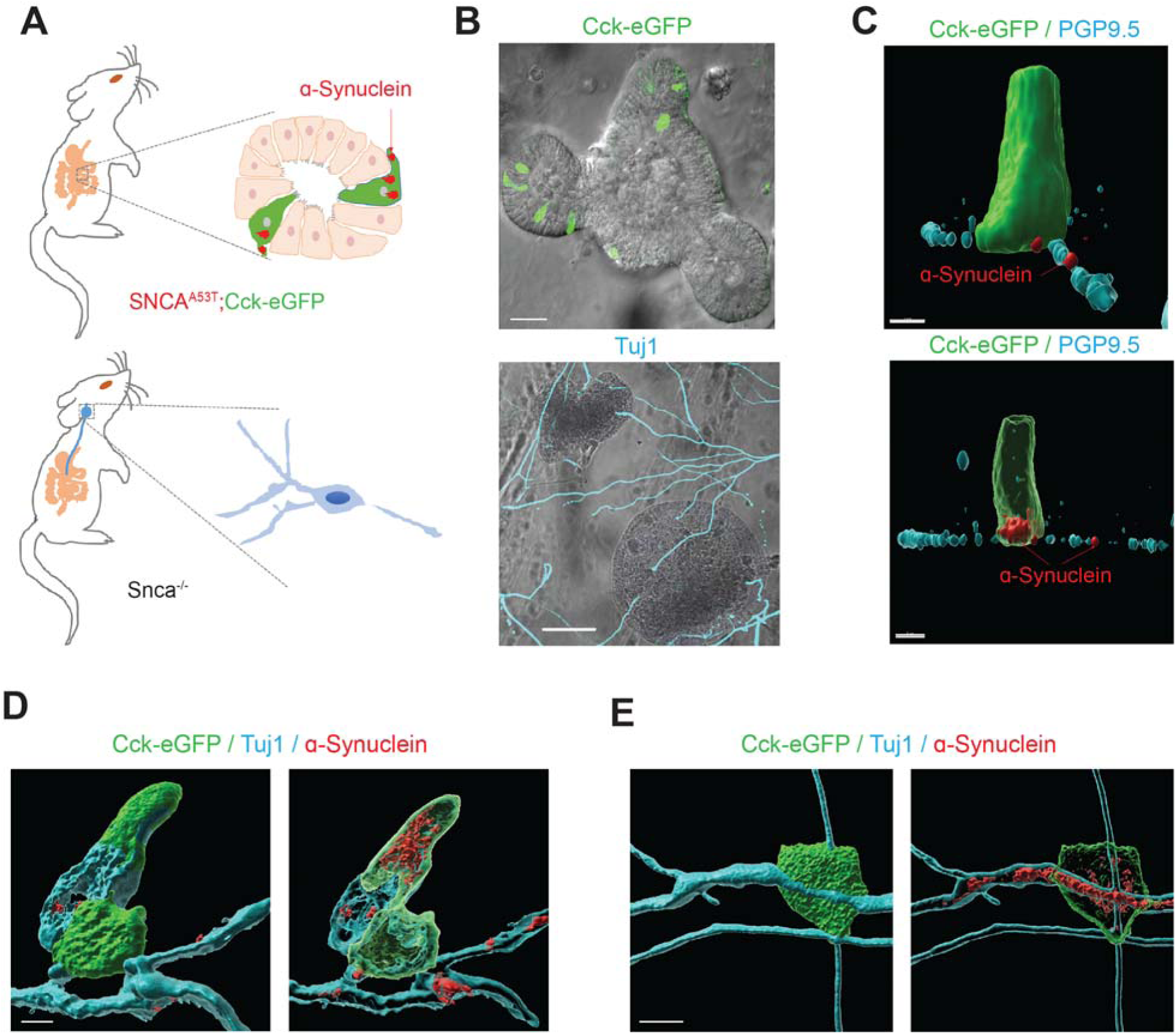
Human A53T α-synuclein protein transfers from gut cells to adjoining vagal neurons. (A) Intestinal organoids were prepared from a SNCA^A53T^ mouse in which CCK-containing cells express enhanced green fluorescent protein (eGFP), and vagal nodose ganglia neurons were isolated from a Snca^-/-^ mouse lacking endogenous α-synuclein. (B) Representative images of organoids and neurons grown in co-culture for five days, with eGFP-positive cells (green) in the organoid, and β-tubulin (cyan) highlighting neuronal processes. (C) Representative high-magnification α-synuclein (red) staining of an eGFP-positive EEC. Red arrow indicates localization to a PGP9.5 (cyan) positive process in an Snca^-/-^ mouse neuron. (D,E) Representative images with neuron-specific class III tubulin (Tuj1, cyan). Surface and (adjacent) intracellular confocal slices are shown. Scale bars are 30 μm for panel B, 3 μm for panel C, and 5 μm for panels D and E.

To determine if gut-to-neuron α-synuclein transfer might occur in intact mammals, and the possible distance to which transferred α-synuclein protein might travel along ganglionic processes (e.g., to the dorsal motor nucleus), we developed a new transgenic strategy to over-express human wildtype and mutant α-synuclein isoforms exclusively in the gut mucosa. This approach is based on villin-Cre conditional expression. Not knowing if aggregation-prone mutated human α-synuclein (i.e., A53T) or wildtype protein may be more likely to transfer from the EECs to nerves, we utilized a recently described “Crainbow” mouse modeling approach to express pathological SNCA gene variants in the same tissue ^52^. We fluorescently barcoded three forms of human SNCA (wildtype, A30P, and A53T mutants) with co-expressed spectrally-resolvable fluorescent proteins from a ROSA-targeting vector for generating α-synuclein-Crainbow mice that we refer to as “SNCAbow” mice (Fig. 3A, Supplemental Fig. 3). SNCAbow mice encode human wildtype α-synuclein co-expressed with nuclear TagBFP (SNCA^WT^:TagBFP), SNCA^A30P^ co-expressed with nuclear mTFP1 (SCNA^A30P^:mTFP1), and SNCA^A53T^ co-expressed with nuclear mKO (SNCA^A53T^:mKO). In the absence of Cre-recombinase activity, only a near-infrared Fluorogen Activating Protein (FAP) is expressed. Crossing SNCAbow mice with villin-Cre (Vil-Cre) mice results in recombination and expression of either SNCA^WT^:TagBFP, SNCA^A30P^:mTFP1, or SNCA^A53T^:mKO (Fig. 3A). From this gene construct, a fluorescent protein and the corresponding α-synuclein were produced as a single mRNA transcript with two distinct polypeptides upon translation [blue TagBFP / SNCA^WT^, turquoise mTFP1 / SNCA^A30P^, and orange mKO / SNCA^A53T^]. We validated expression using intestinal organoids that demonstrate expression of all three fluorescent colors (blue, turquoise, and orange), indicating all three human α-synuclein genes (wildtype, A30P, A53T) were expressed specifically in the intestinal mucosa (Fig. 3B). Interestingly, some cultured organoids expressed only one color, while some organoids expressed multiple colors. This pattern of expression is consistent with clonal growth typical in many organoids where one cell type can dominant others in expansion ^53^. We have previously observed that in the gut mucosa cells, α-synuclein appears exclusively in EECs as observed by immunohistochemistry ^46^. Without addition of external stimulus, pre-formed α-synuclein fibrils, etc., α-synuclein fibril templating activity was detected in the gut organoids in culture nearly comparable to that observed in organoids cultured from the SNCA^A53T^ mice (Fig. 3C,D). In contrast, organoids matched from Snca^-/-^ mice lacked any observable seeding activity.

**Figure 3.**
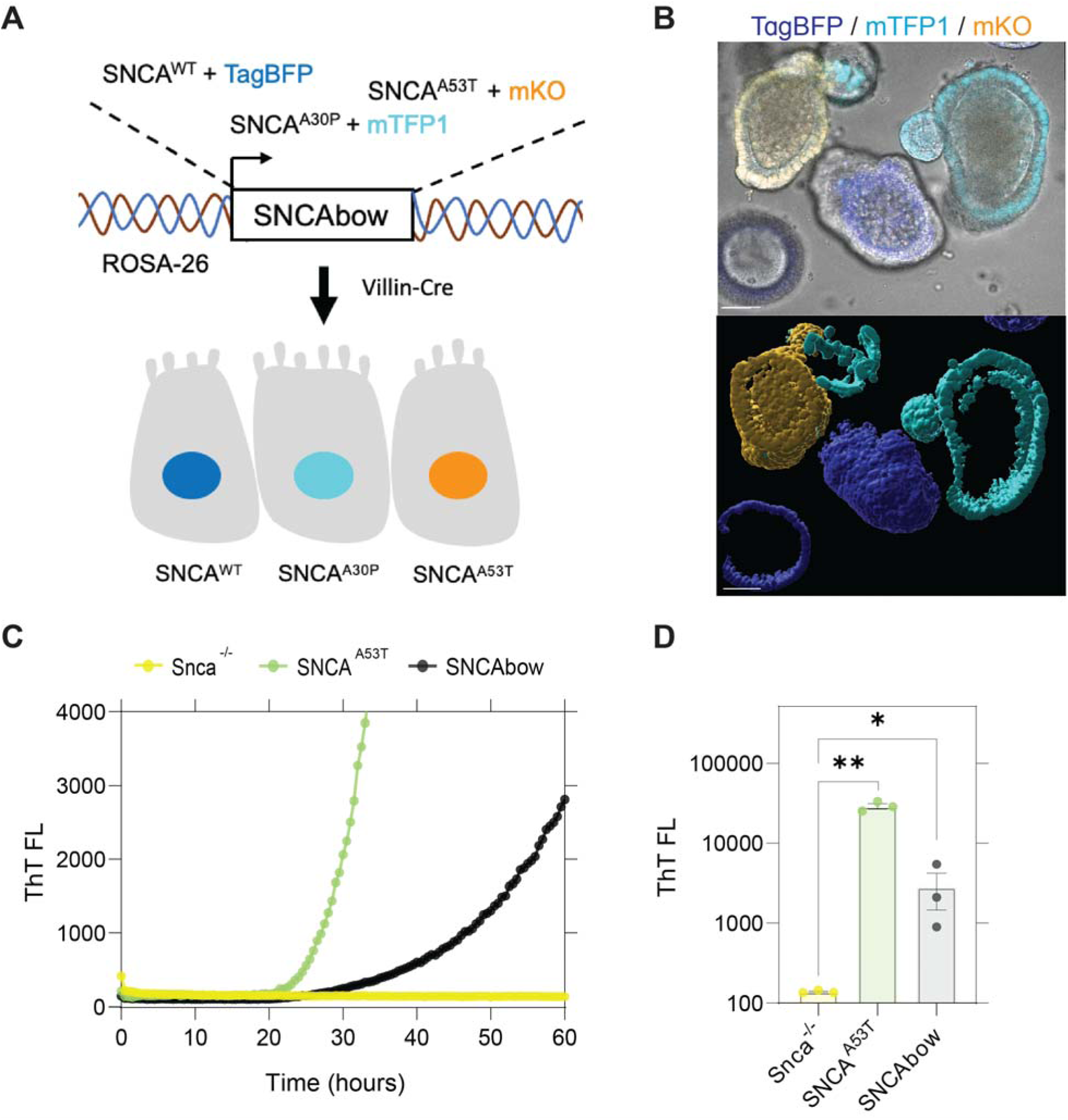
Conditional human α-synuclein expression induces α-synuclein seeding activity in gut organoids. (A) The SNCAbow expression construct contains four tandem cassettes downstream of a chicken-β-actin promoter (not shown). The first cassette (not shown) expresses a chemically inducible near-infrared fluorogen-activating peptide (FAP-Mars1). The next three cassettes encode a unique fluorescent protein (TagBFP:blue, mTFP1:cyan, or mKO:orange) and a corresponding human synuclein protein SNCA^WT^, SNCA^A30P^, SNCA^A53T^. When transgenic mice are mated to the Vil-Cre strain, Cre-mediated recombination by three pairs of orthogonal lox sites (LoxN, Lox2272, LoxP) results in the expression of a single fluorescent protein marker and the corresponding human α-synuclein in any given mucosal cell. (B) Photomicrograph of a small intestine organoids illustrates three fluorescent proteins in the mucosa of a SNCAbow mouse indicating the expression of SNCA^WT^:TagBFP (blue), SNCA^A30P^:mTFP1 (turquoise), SNCA^A53T^:mKO (orange). Scale bar = 30 μm. (C and D) RT-QuIC end-point thioflavin T fluorescence analysis of nodose ganglia from Snca^-/-^, SNCA^A53T^, and SNCAbow mice at 6 months of age. (C) A representative thioflavin T fluorescence profile for these genotypes is provided. (D) End-point values were collected after 100 hours of RT-QuIC relative to negative controls. Data were collected and combined from three mice for each strain in triplicate. Significance was determined by one-way ANOVA with a Dunnett’s post hoc analysis relative to Snca^-/-^, *P<0.05, **P<0.01, n=3.

In the intestine of SNCAbow mice, with α-synuclein protein expression directed by villin-Cre expression, α-synuclein protein was detected in gut mucosal cells via immunofluorescence (Fig. 4A,B) but not in submucosal enteric nerves. We have previously demonstrated that prominent α-synuclein immunostaining in EECs is easily visualized in SNCA^A53T^ mice which contain 4 copies of the human α-synuclein transgene and express at physiological levels in the gut ^46^. However, a sensitive ELISA approach for human α-synuclein protein (that does not cross react with mouse α-synuclein protein) detected the presence of human α-synuclein protein transferred to the vagus of SNCAbow mice (Fig. 4C). Notably, the level of α-synuclein presumably transferred from villin-Cre expressing epithelia (including EECs) in the gut (∼15 pg per total mg of protein) was about one-tenth the level of α-synuclein detected in ganglia in the PAC-human α-synuclein transgenic mouse (Fig. 1C). Despite the lower levels, α-synuclein present in the vagus nerve presumably transferred from gut villin-Cre expressing cells, the RT-QuIC method for measuring fibril-templating activity resolved robust fibril-templating activity in the SNCAbow vagus (Fig. 4D,E). In contrast, no significant templating activity could be identified in non-transgenic mice with only normal concentrations of mouse α-synuclein in the vagus. These results suggest that pathological, templating-positive α-synuclein protein seeds might originate in gut cells in the SNCAbow transgenic mice that transfer α-synuclein protein to the vagus nerve that otherwise lacks intrinsic pathological α-synuclein expression. It was not possible to discern the proportion that each expressed α-synuclein gene (WT, A30P, A53T) contributed to the RT-QuIC signal in the vagus.

**Figure 4.**
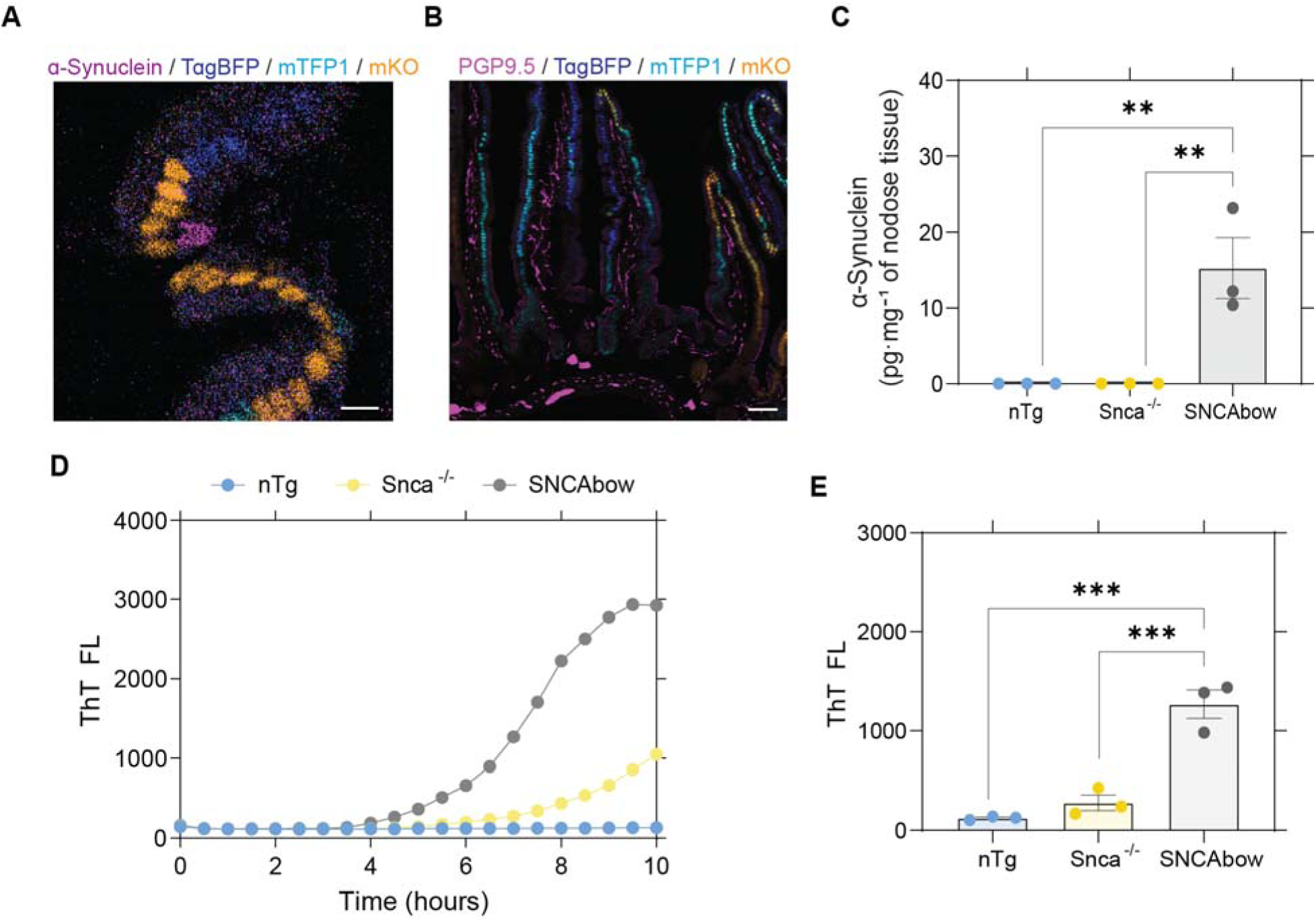
Conditional human α-synuclein expression in gut mucosal cells produces in α-synuclein seeding activity in nodose ganglia. (A) SNCAbow mouse duodenum illustrating expression of fluorescent protein markers BFP (blue), TFP (turquoise), and mKO2 (orange) in mucosal cells (enterocytes and EECs). Alpha-synuclein (magenta) immunostaining is present in a single intestinal mucosal cell consistent with an EEC (Scale bar = 10 μm). (B) Duodenum from SNCAbow mouse illustrating PGP9.5 positive neuronal fibers (magenta) innervating intestinal crypts and villi (Scale bar = 50 μm). (C) ELISA quantification of human α-synuclein protein in nodose ganglia of non-transgenic, Snca^-/-^, and SNCAbow mice. (D) A representative thioflavin T fluorescence profile (RT-QuIC) and end-point analysis of nodose ganglia from non-transgenic (nTg), Snca^-/-^ and SNCAbow mice at 1 month of age. (E) RT-QuIC analysis of nodose ganglia from 6 month old nTg, Snca^-/-^, and SNCAbow mice. Significance was determined by one-way ANOVA with a Dunnett’s post hoc analysis relative to SNCAbow, **P<0.01, ***P<0.001, n=3.

We next tested whether a prophylactic subdiaphragmatic vagotomy procedure might protect the vagus, and ultimately the hindbrain, from the proposed pathological transfer of α-synuclein protein from gut cells. For timed inducible expression of α-synuclein in the gut of the vagotomized adult mice, we crossed the SNCAbow mouse with a tamoxifen-inducible Cre recombinase (SNCAbow;Vil-Cre^ERT2^) to allow aging and subdiaphragmatic vagotomy procedures prior to induction of any human α-synuclein expression (Fig. 5A,B). Three months after tamoxifen treatment, α-synuclein expression was evident in the intestine that was unchanged with vagotomy (Fig. 5C). Fibril templating activity in dissected vagal ganglia was again detected by RT-QuIC after tamoxifen treatment (Fig. 5D,E). In contrast, templating activity was not detected for the mice that underwent the vagotomy procedure, or mice that did not receive tamoxifen treatment. These results suggest that pathological α-synuclein fibril activity likely spreads from the gut cells of the SNCAbow mice to the hindbrain via the vagus nerve (Fig. 5F,G).

## Discussion

This study explores the potential for gut mucosal cells to contribute pathological α-synuclein protein to the nervous system from gut cells. Included are data from three experimental systems: mixed organoid-neuron co-cultures, SNCAbow mice, and inducible SNCAbow mice with or without vagotomy. Collectively, the data indicate that pathological human α-synuclein can transfer from gut mucosal cells to interconnected nerves in mouse models. The main limitations of the study include the reliance on transgenic α-synuclein expression in the gut, as well as a lack of separation of the effects of mutated human pathological α-synucleins from wildtype α-synuclein protein. Future studies may include the creation of new mouse strains that separate different α-synuclein isoforms and expression levels in this process. Further, it remains unclear the molecular nature of the cell-to-cell transfer that might be occurring, whether through bulk exocytosis and uptake ^54–57^, or a more refined local cell-to-cell process like tunneling nanotubes that exist in neuron-to-neuron and glial connections, but have yet to be resolved in gut-to-brain signaling ^58–61^. The organoid-neuron co-culture system provided initial evidence that pathological α-synuclein could transfer from EECs but were limited as an in vitro model. However, confirmation that α-synuclein spread also occurs in vivo in SNCAbow mice supports the relevance of the co-culture system which could be used to explore the cellular mechanisms of EEC-to-neuron transfer. Nevertheless, these results highlight the existence of a new non-neuronal potential source for pathological α-synuclein protein that may contribute to the pool of abnormal α-synuclein at some point in disease.

**Figure 5.**
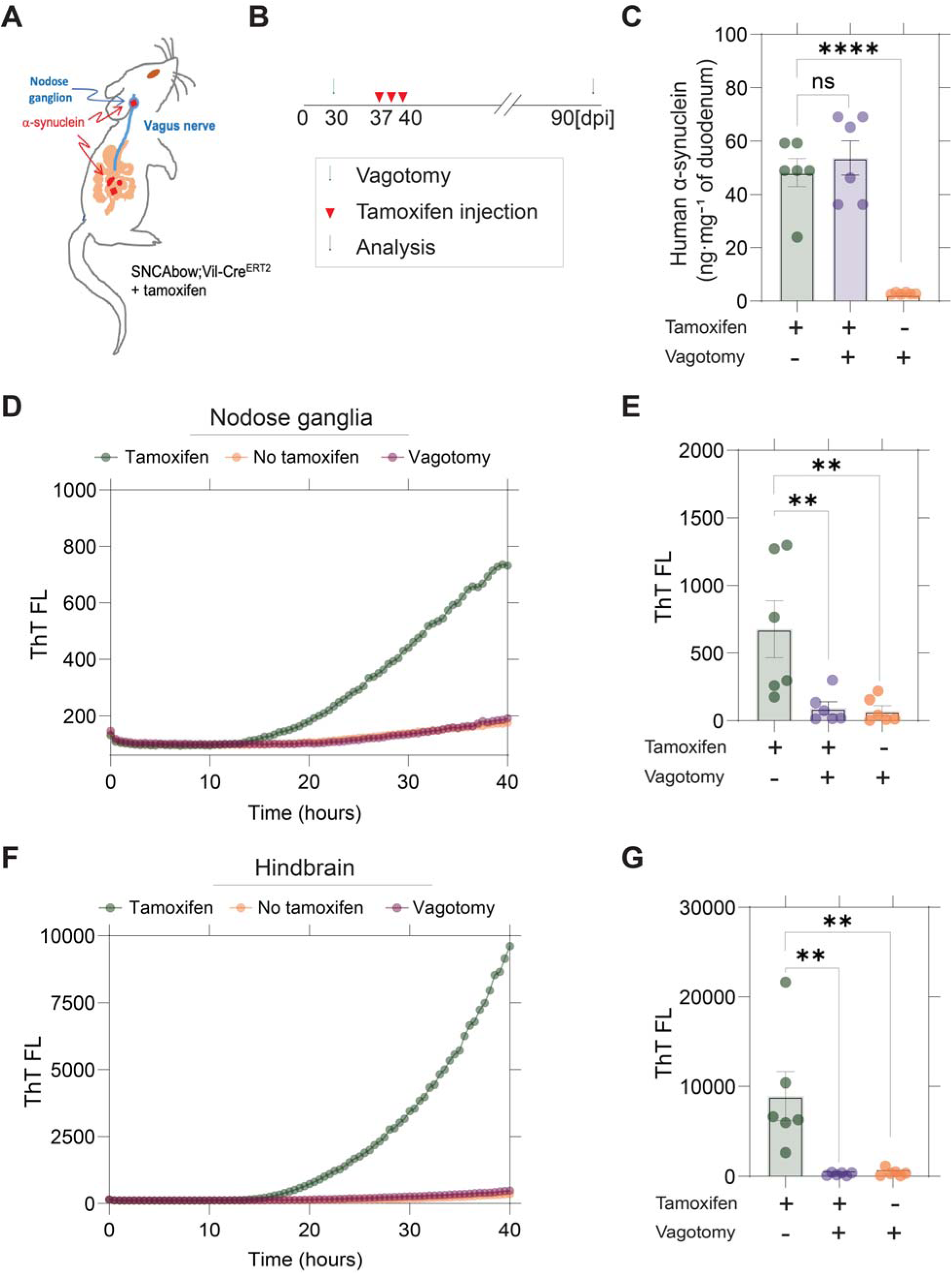
Vagotomy spares the nodose ganglia from α-synuclein seeding activity and prevents spread to the hindbrain. (A and B) Experimental model. SNCAbow;Vil-Cre^ERT2^ mice underwent bilateral sub-diaphragmatic vagotomy or sham surgery 1 week before tamoxifen treatment. (C) ELISA measurements of α-synuclein protein in the gut three months after tamoxifen treatment. RT-QuIC analysis of (D,E) vagal nodose ganglia and (F,G) hindbrain analyzed three months after tamoxifen treatment. Representative thioflavin T fluorescence profiles are shown in D and F. Data points in B, E, G represent the Significance was determined by a one-way ANOVA with a Dunnett’s post hoc analysis relative to tamoxifen treated and no vagotomy groups. **P<0.01, ****P<0.0001, n=6. d in the nodose ganglion. Bar = 20 μm in all images.

Braak highlighted the hypothesis that PD pathology may arise in peripheral nerves and spread to the central nervous system in some cases ^22^. The identification of α-synuclein aggregates in enteric nerves, before their appearance in the brain, may be consistent with gut-to-brain pathological spread ^29,62^. However, according to prion-like activity associated with α-synuclein in disease (especially in models of disease), and α-synuclein cell-to-cell transfer observed in different paradigms, corrupted and misfolded α-synuclein need not originate in the cells that wind up with large observable aggregates. Based on the evidence presented herein, we hypothesize that the gut mucosa, especially EECs, may contribute pathological misfolded α-synuclein to vulnerable efferent and afferent projections of the vagus nerve, which might predispose to the risk of Lewy body disease. Notably, EECs in the mucosa make physical contact with both the microbiome and ingested toxicants like pesticides in the gut lumen, as well as contact nerve fibers on the other side, providing a ripe opportunity to better understand the pathobiology of the microbiome and toxin exposures in PD vulnerability.

Vagotomy is known to abolish the spread of gut-injected recombinant preformed fibrils (PFFs) to the brainstem, indicating that the transmission mechanism might occur via the vagus nerve ^25^. Herein, pathological α-synuclein was introduced not by the injection of pre-formed α-synuclein fibrils, but through the over-expression of either wildtype of mutated human α-synuclein isoforms in gut mucosal cells. According to ELISA analysis, the over-expression achieved was modest and in line with PAC-transgenic expression that has been described as physiologically comparable to α-synuclein expression in humans ^32^. Although EECs are distributed throughout the gastrointestinal tract, we focused on the proximal small intestine where vagal innervation is most abundant.

Due to their high sensitivity, seeding amplification assays have been used with increasing frequency to detect early pathology in PD and distinguish PD from non-affected individuals ^63,64^. We used RT-QuIC to detect the early spread of α-synuclein to obtain a semi-quantitative estimate of α-synuclein abundance in tissue lysates. This approach is similar to human seeding assay results for tissues like duodenum and CSF 20,65.

Transplants of gut microbiota from PD patients into gnotobiotic mice accelerate α-synuclein pathology and PD-associated behavioral changes in mice bearing human α-synuclein ^66^. The mechanism of formation of pathological α-synuclein in response to the imbalanced gut microbiota remains elusive, albeit we hypothesize here the role of EECs as a possible target given that they are directly exposed and respond to microbes and microbial metabolites ^67,68^. Recently it has been demonstrated that Akkermansia muciniphila, a bacterial strain residing in the gastrointestinal tract and associated with PD, led to α-synuclein aggregation in an enteroendocrine cell line ^69^. Therefore, although the specific habitat for pathological α-synuclein in the gut is unknown, our identification of α-synuclein in EECs and the location of EECs at the interface between the gut lumen, rich with microbiota and enteric nerve fibers, has raised the possibility that EECs may be a source for the formation and possible spread of pathological α-synuclein ^46^. Alternatively, pathological α-synuclein from the brain may spread into EECs that might then transfer to interconnected neurons otherwise devoid of pathological α-synuclein. The initial step in determining whether EECs are capable of transmitting pathological α-synuclein was confirmed by our in vitro studies demonstrating the transfer of human A53T α-synuclein from EECs in organoid culture onto isolated Snca^-/-^ nodose ganglion neurons. Using nerves lacking endogenous α-synuclein together with an antibody specific for human α-synuclein, it was possible to establish that EECs were the source of the α-synuclein immunoreactivity appearing in the nearby neurons. This experiment demonstrated that EECs possess the ability to transfer endogenous α-synuclein onto adjacent nerves, a process that could be the initial step in the spread of pathological α-synuclein into the nervous system in some patients in disease. Despite the direct transfer of α-synuclein from EECs to nodose neurons in vitro, whether other cell types contribute to the spread in vivo is unknown. EECs come into contact with glia ^45^ and connect to enteric neurons PMID: 25555217 as well as the vagus nerve ^41^. It is possible that α-synuclein can spread to enteric glia ^70^ and thus, provide a potential route for the spread of α-synuclein. It remains to be determined what role glia might have in the spread of α-synuclein to enteric nerves including the vagus that extends beyond the gut.

EECs are sensory cells of the intestine that communicate directly with afferent vagal nerve fibers ^41^. The function of α-synuclein expression in these cells remains to be clarified. The close proximity of EECs to the nerve processes prompted us to look for α-synuclein in the vagus in the gut-restricted models. Our detection of α-synuclein seeding activity in the vagus in the SNCAbow mice is consistent with reports of neuropathology in the sensory branch of the vagus nerve in patients with PD ^71^ and the possible spread of PFFs to the nodose ganglia in experimental models of PD ^27^. However, it is not known if this is a common route for the spread of PD.

It is generally believed that templating of α-synuclein in the recipient cell is necessary to facilitate cell-to-cell propagation of pathological α-synuclein. It is notable in this regard that α-synuclein protein does not appear to reside in the dendrites of vagal afferents where it would be available for such templating ^72^. Nevertheless, the in vitro co-culture experiments demonstrated that α-synuclein in the recipient neuron is not required for α-synuclein uptake. The appearance of α-synuclein templating activity in SNCAbow nodose ganglia in vivo suggest that the transgenic human α-synuclein can spread at least as far as the vagal ganglia. Therefore, it is plausible that α-synuclein may transfer to vagal afferents and spread intracellularly before any templating with endogenously expressed protein occurs in the axonal compartment of the neuron. The question of whether transmission occurring along vagal afferents, or descending back to the gut from the brain, preferentially drives disease phenotypes in disease is a major question that can be tackled in-part in the future with additional conditionally-restricted α-synuclein expression models.

## Methods

### Mice

All experiments were performed with approval by the Duke University Institutional Animal Care and Use Committee. Mice expressing enhanced green fluorescent protein in CCK cells [Tg(Cck-EGFP)BJ203Gsat], referred to as Cck-eGFP, were obtained from Mutant Mouse Resource and Research Center (RRID:MMRRC_000249-MU), Missouri and the colonies were maintained on a Swiss Webster background (Taconic Biosciences) ^73^. FVB;129S6-Snca1Nbm Tg(SNCA*A53T)1Nbm Tg(SNCA*A53T)2Nbm/J mice (PAC-SNCA^A53T^) ^32^ and Snca^-/-^ mice ^74^ were obtained from Robert L. Nussbaum (RRID:IMSR_JAX:010788). The generation of SNCA^A53T^; Cck-eGFP mice has been described previously ^46^ and are referred to herein as SNCA^A53T^.

Human wildtype, A53T or A30P forms of α-synuclein were expressed from a modified brainbow gene construct that also harbored three fluorescent proteins (blue, turquoise, and orange). In this manner only one fluorescent protein could be expressed from each copy of the construct. Cre recombinase is driven by the villin promoter (Vil-Cre) targeted expression to mucosal cells of the gastrointestinal tract with different fluorescent proteins expressing in each stem cell. SNCAbow mice were generated using a technique that was recently described ^52^. Briefly, C-terminal tagged α-synuclein genes (SNCA tagged with V5, hSNCA-A30P mutant tagged with 3XHA, and SNCA-A53T mutant tagged with Myc) were amplified by PCR and pENTR™ plasmids were generated. These plasmids were sequenced in entirety at Massachusetts General Hospital Center for Computational & Integrative Biology DNA Core (https://dnacore.mgh.harvard.edu/new-cgi-bin/listing.action) and cloned by Infusion cloning into a ROSA26 mouse targeting vector adapted for Gateway cloning. Plasmid DNA harvested from bacterial colonies was mapped by restriction enzyme digestion analysis and positive colonies were identified. The entire plasmid was sequenced, linearized with XhoI, and transfected into G4 ES cells (129/B6N hybrid ES line, (MMRRC Cat# MMRRC:011986-UCD, RRID:CVCL_E222). Putative positive ES cell clones were processed and validated across the homology arms by PCR. DNA from two selected colonies was amplified using LA Taq DNA polymerase (TaKaRa) and the region between the homology arms was sequenced. ES cells from the selected clone was microinjected into ICR/Hsd morulae to produce chimeric mice. The transgenic mouse was mated with ROSA FLPe [Jax-129S4/SvJaeSor-Gt(ROSA)26Sortm1(FLP1)Dym/J (Jackson laboratory, RRID:IMSR_JAX:003946)] to excise the neomycin cassette, and then mated with Vil-Cre mouse (Jax-B6.Cg-Tg(Vil1-cre)997Gum/J ((Jackson laboratory, RRID:IMSR_JAX:004586)) or Villin-Cre^ERT2^ (kind gift of Sylvie Robine, RRID:IMSR_GPT:T004829) for expression of fluorescent proteins and associated α-synuclein transgenes in intestinal mucosal cells.

### Vagotomy and tamoxifen treatment

Surgical subdiaphragmatic vagotomy was performed in 1 month old male and female SNCAbow;Vil-Cre^ERT2^ mice . Mice were anesthetized with ketamine (50-100 mg per kg) and an abdominal laparotomy was performed. Immediately below the diaphragm, the vagus nerve was identified and isolated from surrounding connective tissue and vessels. A 2 mm section of the vagus nerve was excised and the surgical wound was closed with surgical clips. Mice were administered analgesics and observed daily for 5 days for any signs of distress . In ‘sham’ operated animals, abdominal laparotomy was performed, and the vagus nerve was exposed but not excised. Weight loss of ∼15% was noted in mice undergoing vagotomy compared to ‘sham’ surgery. One week following the surgery mice were treated with tamoxifen (5 mg per kg) or vehicle administered by intraperitoneal injection daily for five days.

### Preparation and culture of mouse organoids

Mouse small intestine was dissected, gently flushed with ice cold phosphate buffered saline (PBS) (pH 7.4)/Primocin (1:1000) (InvivoGen, Cat# ant-pm-1) and cut into ∼0.5 cm pieces that were placed in 7.5 mL cold PBS/EDTA (3 mM)/Primocin/Y27632 (1:1000) (ApexBio, Cat#A3008-200) containing penicillin-streptomycin (Gibco, Cat#15140-122) and gently shaken for 15 min at 4°C. The intestinal tissue was transferred to fresh EDTA/PBS/Primocin/Y27632, shaken for 25 min at 4°C, and transferred to PBS. Tissue was then transferred to PBS/Y27632, shaken for 2 min, and filtered through 70 μm mesh, examined under a microscope, and aliquoted at a density of 50 crypts in 15 μL growth factor reduced Matrigel (Corning, Cat#354230). The suspension was centrifuged at 1500 rpm for 5 minutes at 4°C and the pellet was resuspended in cold growth factor reduced Matrigel and aliquoted (15 μL per well) in a 48 well plate (Eppendorf, Cat#0030723113). Matrigel was let to polymerize for 30 minutes at 37°C. To each well, 200 μL of pre-warmed (37°C) Intesticult Media (Stem Cell Technologies, Cat#06005) containing Primocin. Media were changed every two days and organoids were split weekly.

### Isolation and co-culture of nodose ganglion neurons

Nodose ganglia were dissected from Snca^-/-^ mice and placed in 300 μL ice cold mouse Intesticult media (StemCell Technologies, Cat#06005) containing nerve growth factor-2 (NGF) (25 ng per mL, Sigma, Cat#N6009) and Liberase (0.156 mg per mL). After incubating at 37°C for 30 minutes, the supernatant was replaced with 500 μL Intesticult media containing NGF. Tissue was dissociated by pipetting, filtered through a 70 μm mesh, and centrifuged at 800 rpm for four minutes. The pellet was resuspended in fresh media, mixed with growth factor reduced Matrigel (Corning, Cat#354230) and added to intestinal organoid cultures (at least four weeks old). The nodose ganglia/organoid mixture was incubated in an 8-well chamber slide at 37°C for 30 minutes to allow polymerization. Subsequently, pre-warmed Intesticult media containing NGF was added and the cell mixture were grown for an additional 5-8 days prior to imaging.

### Immunostaining of organoids

Whole mount organoid staining was performed as described previously ^75^ with slight modifications. After removal of media, organoids were fixed in 4% paraformaldehyde in PBS, (pre-warmed at 37°C to prevent Matrigel depolymerization) for 20 minutes at room temperature. Organoids were permeabilized with pre-warmed 0.5% Triton X-100 in PBS, followed by three washes with 100 mM glycine (Invitrogen, Cat# 10977-023). After blocking with 5% bovine serum albumin/5% donkey serum/PBS for 2 hours at room temperature, primary antibody was added and incubation continued in a humidified chamber for 16 hours at 4°C. Primary antibodies used for immunostaining included: Rabbit CCK ^73^, rabbit α-synuclein (Abcam Cat# ab138501, RRID:AB_2537217, at 1:1000), guinea pig PGP9.5 (Abcam Cat# ab10410, RRID:AB_297150, at 1:100), chick beta-tubulin III, Tuj1 (Neuromics Cat# CH23005, RRID:AB_2210684, at 1:100), and chick GFP (Abcam Cat# ab13970, RRID:AB_300798, at 1:1000). Slides were washed three times for 20 minutes each in IF buffer (PBS containing 0.1%BSA, 0.2% Triton X-100, 0.05% TWEEN 20). Secondary antibodies (in IF) were applied for 1 hour at room temperature in the dark. Secondary antibodies included: Donkey anti-chicken Alexa Fluor 488 (Jackson ImmunoResearch Labs Cat# 703-545-155, RRID:AB_234037, at 1:500), and Donkey anti-mouse Alexa Fluor 568 (1:500) and, Donkey anti-guinea pig Alexa Fluor 647 (Jackson ImmunoResearch Labs Cat# 706-605-148, RRID:AB_2340476, at 1:250). Slides were washed three times, 20 minutes each wash, with IF buffer. DAPI was applied for 5 minutes at room temperature, washed in PBS, and mounted with ProLong Gold (ThermoFisher, Cat#P36930).

### Immunofluorescence of duodenal and nodose tissue sections

To characterize expression of transgenes and fluorescent proteins, mice were anesthetized with a mixture of xylazine and ketamine and perfused with ice cold 3.5% freshly depolymerized paraformaldehyde. Intestinal tissue was harvested, post-fixed, and cryopreserved in graded sucrose solutions. The tissue was embedded in OCT and cryosections (10 – 20 μm thickness) were collected on plus charged slides. Immunostaining was performed as described previously ^46^. The following primary antibodies were used: chick GFP (Abcam Cat# ab13970, RRID:AB_300798, at 1:1000), rabbit α-synuclein (Abcam Cat# ab138501, RRID:AB_2537217, at 1:1000), sheep α-synuclein (Abcam Cat# ab6162, RRID:AB_2192805, at 1:1000), rabbit PGP9.5 (Millipore Cat# AB1761-I, RRID:AB_2868444, at 1:50).

### α-Synuclein protein expression and purification

Plasmid construct (pRK172-human-lil-synuclein) encoding WT-human-α-synuclein was expressed in BL21-CodonPlus (DE3-RIL) cells (Agilent, Cat# 230245-41). Protein expression was induced with 0.1 mM IPTG at cell density OD (600 nm) 0.8 and overnight incubation at 18°C with continuous shaking. Cell pellets were lysed in 0.75 M NaCl, 10 mM Tris HCl, pH 7.6, 1 mM EDTA, 1 mM PMSF and sonicated at 30% power (Fisher 500 Dismembrator) for 1 minute followed by boiling the cell suspension 15 minutes. Centrifuged and filtered samples were dialyzed against 10 mM Tris HCl, pH 7.6 with 50 mM NaCl, 1 mM EDTA, 1 mM PMSF. The suspension was passed through a HiPrep Q HP 16/10 Column, 1 x 20 mL (GE Healthcare) on an ÄKTA pure protein purification system (Cytiva, formerly GE Healthcare) with running buffer composed of 10 mM Tris pH 7.6, 25 mM NaCl, and eluted with a linear gradient application of high-salt buffer (10 mM Tris HCl, pH 7.6, 1 M NaCl). Fractions containing a single-band of α-synuclein were identified by Coomassie staining of SDS-PAGE gels and were dialyzed and concentrated. Purified monomer protein was subjected to two to three rounds of endotoxin removal (Endotoxin removal kit, GenScript) until a level of <0.1 EU per mg was achieved. Endotoxin levels were determined using a LAL chromogenic endotoxin quantification kit (GenScript).

### Preparation of mouse tissue homogenates for RT-QuIC assay

Mouse tissues (10% weight per volume) were prepared by homogenizing the tissue in PBS supplemented with 1% Triton X-100 followed by probe tip sonication for 1 min (5 sec On, 15 sec Off) at 10% amplitude (Fisher 500 Dismembrator). Sonicated samples were centrifuged for 20 min at 20,000 x g at 4°C and supernatants were aliquoted for further applications. For RT-QuIC analysis, 1% of homogenates were added into the reaction.

### Real time-quaking induced conversion assay (RT-QuIC)

RT-QuIC analysis was performed in clear bottom, ultra-low binding 384-well plates (Corning) loaded with one zirconia silica bead (2.3 mm, OPS Diagnostics) per well. Reaction conditions included 10 μM Thioflavin T (ThT) in sterile PBS, 20 μM of human α-synuclein monomer and 10% of tissue homogenates (1% weight per volume). Human α-synuclein monomer was passed through an ultra-0.5 centrifugal filtration system (100 kDa MWCO, Amicon) to remove high-molecular-weight aggregates. Reactions were performed in triplicate and each technical and biological experiment was repeated three times. Standard curves with serial dilutions of recombinant α-synuclein fibrils in 1% of matrix homogenate from Snca^-/-^ mice were included for each reaction in order to confirm the efficiency, sensitivity and specificity of the reaction (Supp. Fig.1). Serial dilutions of recombinant fibrils were prepared according to approximate molecular weight calculated via Dynamic Light Scaterring measurements and estimated structural arragments as described ^47^. Negative control samples including the lysates from Snca^-/-^ tissues were used to exclude any background signals likely occurring due to tissue origins. The RT-QuIC assay was conducted using a OMEGA BMG plate reader for 15-90 hours with 60 sec of shaking at 700 rpm and 60 sec of rest. The ThT signal was monitored every 30 min at 448-10 excitation and 482-10 emission.

### Chemiluminescence-enhanced ELISA measurements for human-α-synuclein

Determination of human α-synuclein concentration in nodose ganglia was performed according to manufacter’s instructions (Biolegend, Cat#844101). The 1:10 diluted samples were run in triplicates and reference standard curve ranged from 6.1 – 1500 pg per mL. Luminescent signal was recorded on a ClarioStar plate reader (BMG).

## Authors’ contributions

RC and AS contributed to the experimental design, conducted experiments, and wrote the manuscript. KRC, SK, and SMS conducted experiments and reviewed the manuscript. JCS contributed to the experimental design and reviewed the manuscript. ABW contributed to the experimental design and wrote the manuscript. RAL designed the experiments and wrote the manuscript.

## Acknowledgments

This work was supported by NIH grant R01 DK-120555, Department of Veterans Affairs grant I01 BX002230, and the Holland-Trice Award from Duke University to R.A.L; NIH grants R01 NS-064934 and P50 NS-108675 to A.B.W; and 5K22 CA-212058-03 to J.C.S. This research was funded in part by Aligning Science Across Parkinson’s award ASAP-020527 through the Michael J. Fox Foundation for Parkinson’s Research (MJFF). For the purpose of open access, the authors have applied a CC BY public copyright license to all Author Accepted Manuscripts arising from this submission. The authors acknowledge the support of Duke University’s Light Microscope Core Facility, Transgenic Mouse Facility, and the Duke Cancer Institute.

The authors thank Senthil Karuppagounder for technical support.

