## Supplemental Figures for "Gut mucosal cells transfer α-synuclein to the vagus nerve"

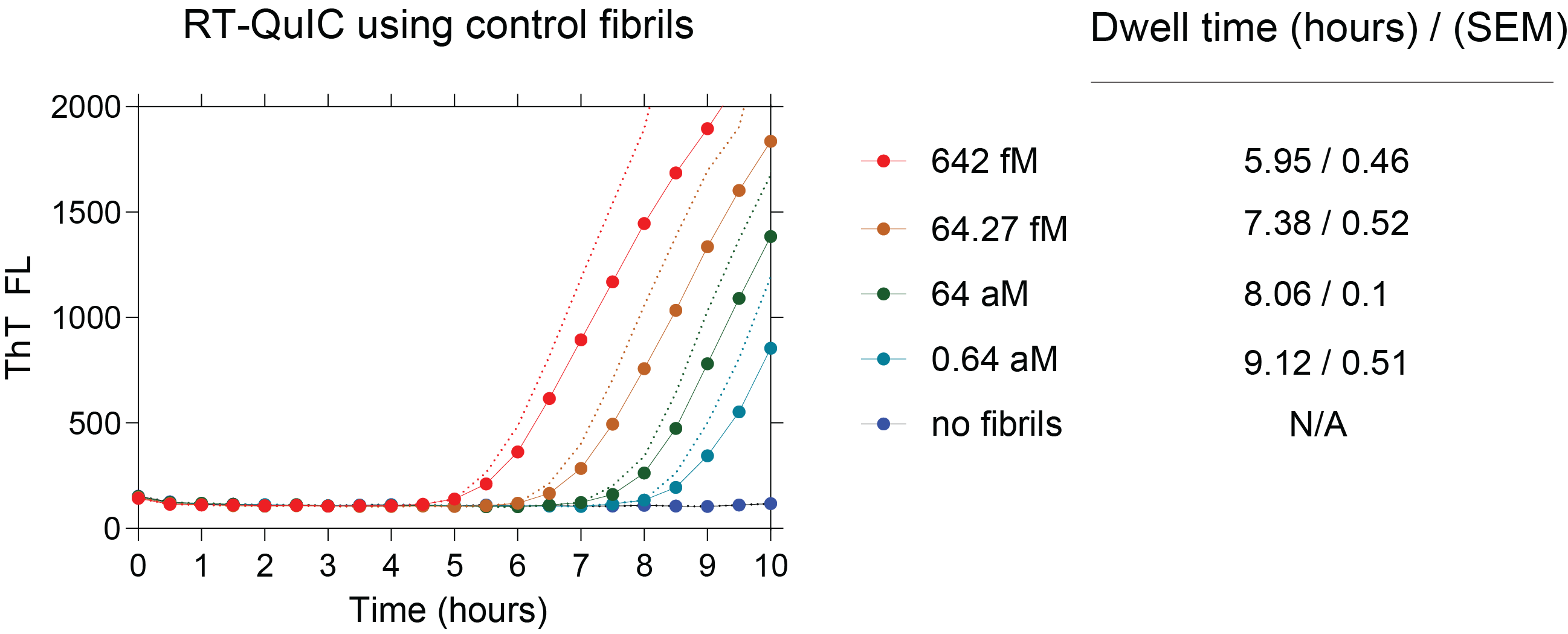

**Supplemental Figure 1. Spike-in recombinant fibril seeds demonstrates sensitivity of RT-QuIC assays in protein lysates from *Snca* knockout nodose ganglia lysates**

Histograms showing a titration of concentrations of fibril seeds over four orders of magnitude in RT-QuIC reactions. Reaction buffer includes 1% of tissue lysates from nodose ganglia from *Snca^-/-^* mice to account for matrix effects on α-synuclein aggregation. Dwell times are with respect to increases in ThT 2xS.D. above background (e.g., no fibril) parallel runs. Each dot shows the mean value from triplicate independent reactions (SEM indicated with dashed lines).

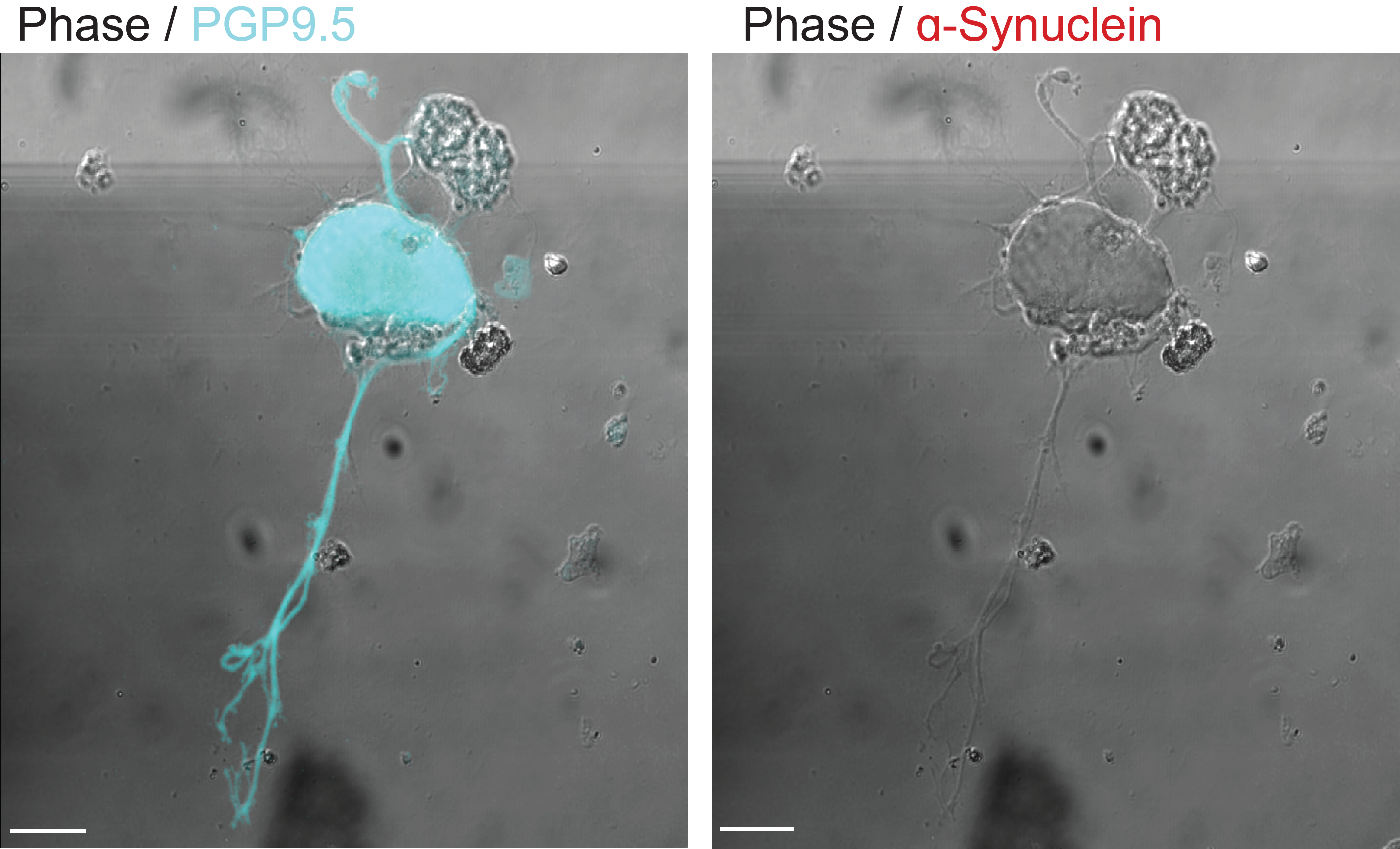

**Supplemental Figure 2.** **Nodose ganglion neuron from a *Snca^-/-^* mouse lacks** α**-synuclein.** (Left panel) Confocal photomicrograph of a nodose ganglion neuron isolated from a *Snca^-/-^* mouse grown in culture for one week and stained with an antisera specific for the pan-neuronal marker PGP9.5 (turquoise). (Right panel) Immunofluorescence staining with antisera for α-synuclein (red) (Abcam, ab138501, MJFR1). As expected, neurons from *Snca^-/-^* mice have no immunodetectable α-synuclein. Bar = 30 μm.

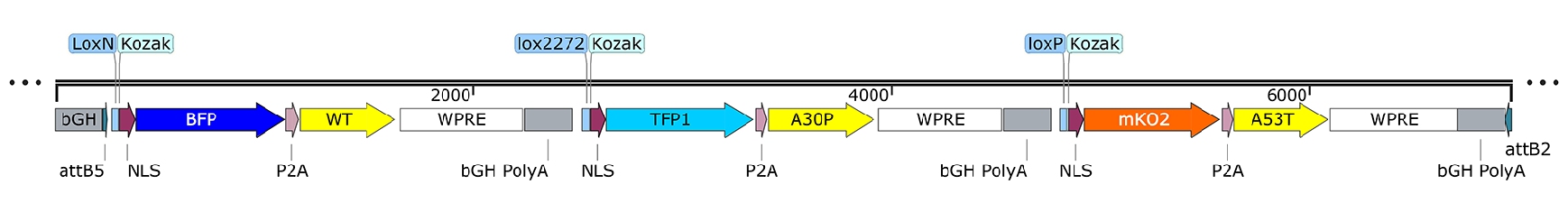

**Supplemental Figure 3.** **SNCAbow construct expresses three human α-synuclein** **proteins.** (A) The SNCAbow expression construct contains four tandem cassettes downstream of a chicken-β-actin promoter (not shown). The first cassette (not shown) expresses a chemically inducible near-infrared fluorogen-activating peptide (FAP-Mars1). The next three cassettes encode a unique fluorescent protein (TagBFP:blue, mTFP1:turquiose, or mKO:orange) and a corresponding human α-synuclein protein SCNA^WT^, SCNA^A30P^, SCNA^A53T^. When transgenic mice are crossed with mice expressing Cre-recombinase, Cre-mediated recombination by three pairs of orthogonal lox sites (LoxN, Lox2272, LoxP) results in the expression of a single fluorescent protein marker and the corresponding human α-synuclein in any given mucosal cell.

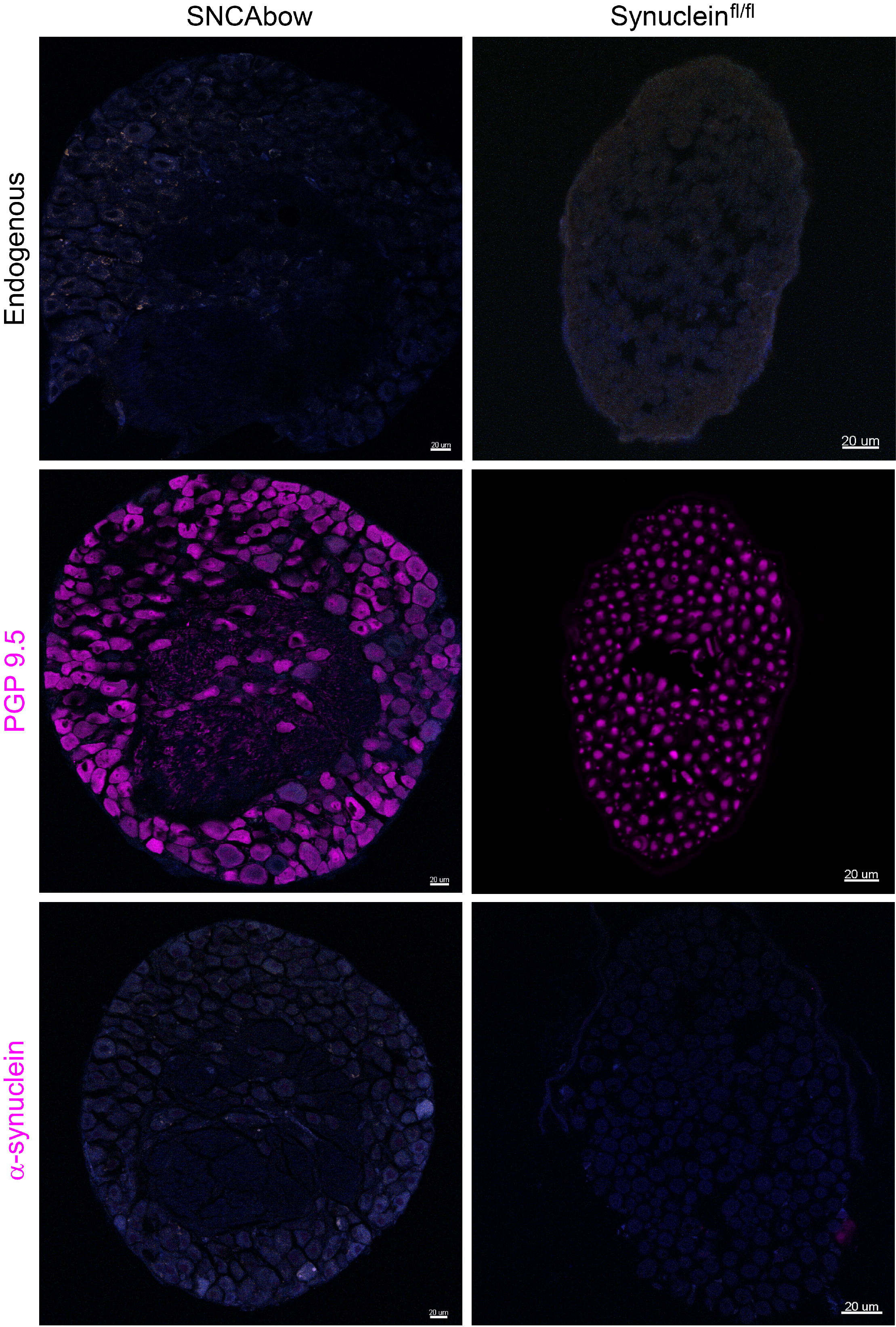

**Supplemental Figure 4.** **Nodose ganglia of SNCAbow mice do not express aberrant transgene fluorescence.** Confocal images showing endogenous fluorescence (no immunostaining) (top row), PGP9.5 immunostaining (middle row), and α-synuclein immunostaining (bottom row) in the nodose ganglion of SNCAbow and α-synuclein^fl/fl^ mice (in the absence of Vil-Cre). Unlike the duodenum, endogenous fluorescence was not detected in the nodose ganglion of SNCAbow mice. PGP9.5 immunostaining is present in nerve bundles of both genotypes of mice. α-Synuclein (Abcam, ab138501, MJFR1) could not be detecte
